## Supplementary figures and tables for "Artificial selection improves pollutant degradation by bacterial communities"

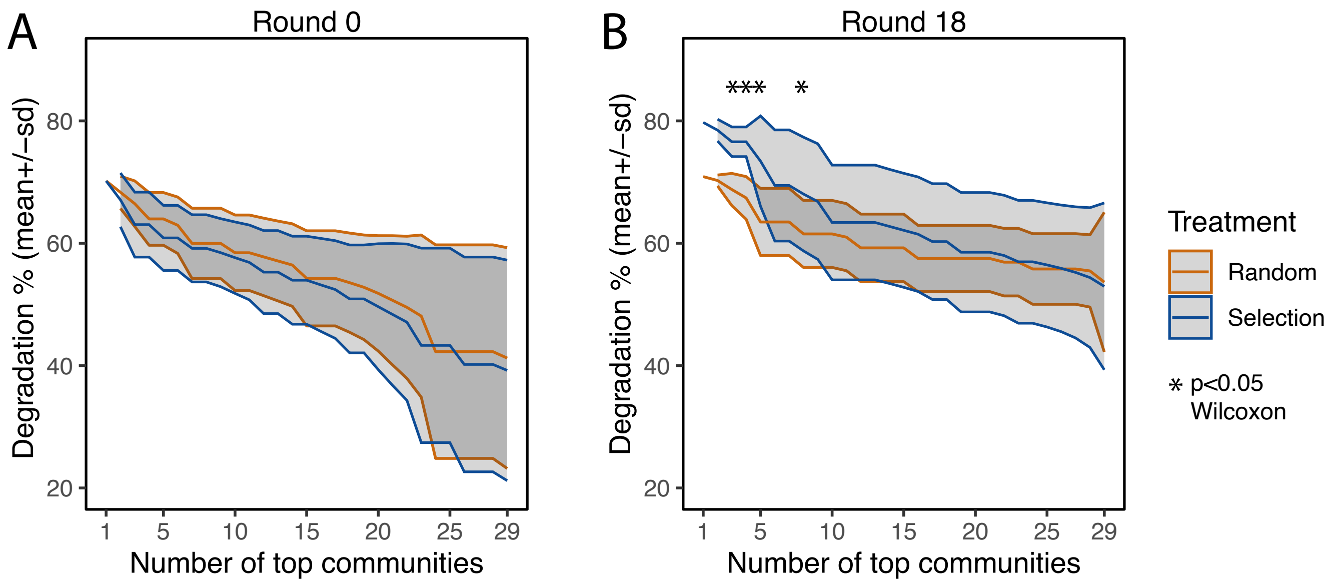


**Fig. S1.** Justification for why we chose to compare the top 5 communities between treatments. Both plots show the mean±std (central and shaded areas) of the degradation percent of the top x communities in round 0 (A) and round 18 (B). Naturally, the fewer communities we take, the smaller the standard deviation becomes. While in round 0 there is no difference between the treatments, at round 18, the selection treatment becomes significantly better (Wilcoxon rank sum exact test, $p < 0.05$) only when we take the top few communities, indicated with asterisks. The noise is uninteresting from the perspective of an optimization algorithm, as what we are striving toward is to find the very best community. However, this analysis does suggest that we should work to reduce the noise, for example by lowering the fraction of communities receiving migrants.

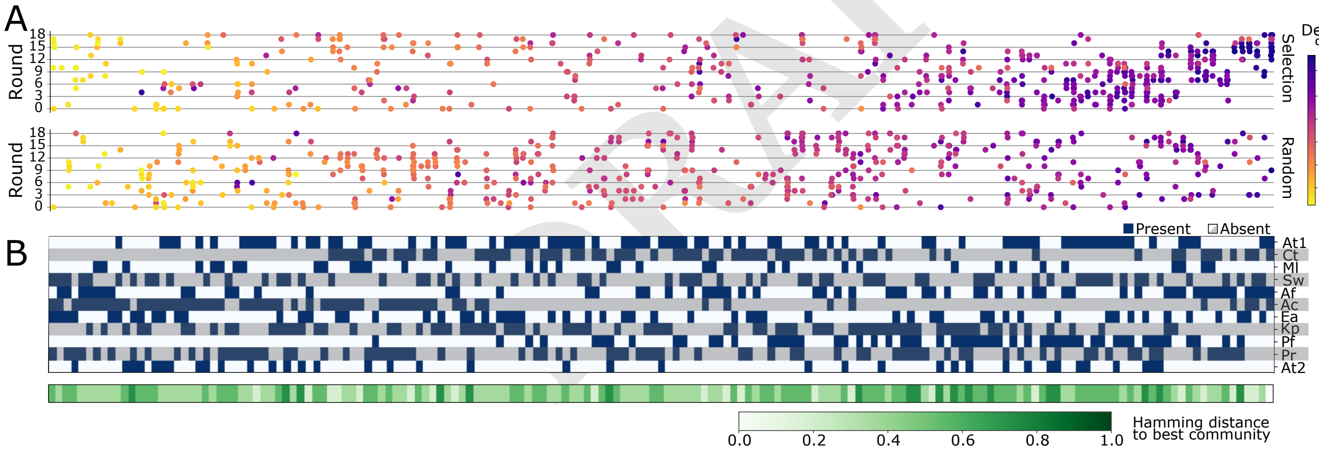


**Fig. S2.** A) Community composition (x-axis) vs. degradation score (hue, color bar) for all 167 tested communities over the 18 rounds of selection (y-axis) in the selection (top panel) and random (bottom panel) treatments. The x-axis is ordered by increasing degradation scores (averaged over all instances of the same species composition). Note that these are degradation percentages, not final community scores (extinctions not considered). B) Community composition corresponding to panel A, showing the presence (dark blue) or absence (white/grey) of each species, and illustrating the difference in composition by the Hamming distance (i.e. the number of substitutions needed to transform a given community to another) to the community with the highest degradation score at the bottom. This figure overlaps with Fig. 1C, D in the main text, but includes all 167 tested communities as opposed to only the best third (56) of all communities tested.

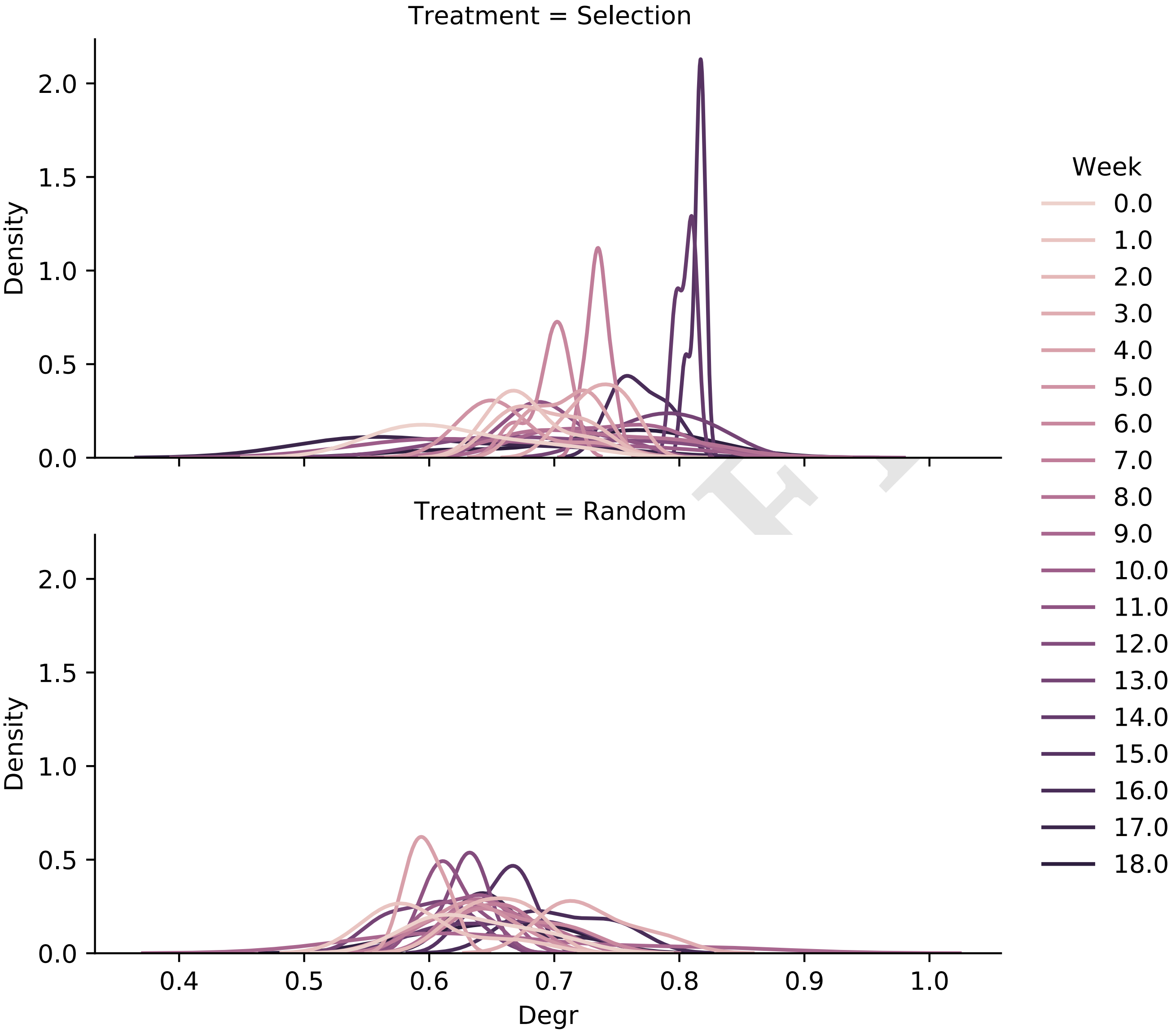


**Fig. S3.** Distribution (Gaussian kernel density estimates) of degradation scores (x-axis, "Degr") of all 30 communities, colored by week. Top row: selection treatment, bottom row: random control.

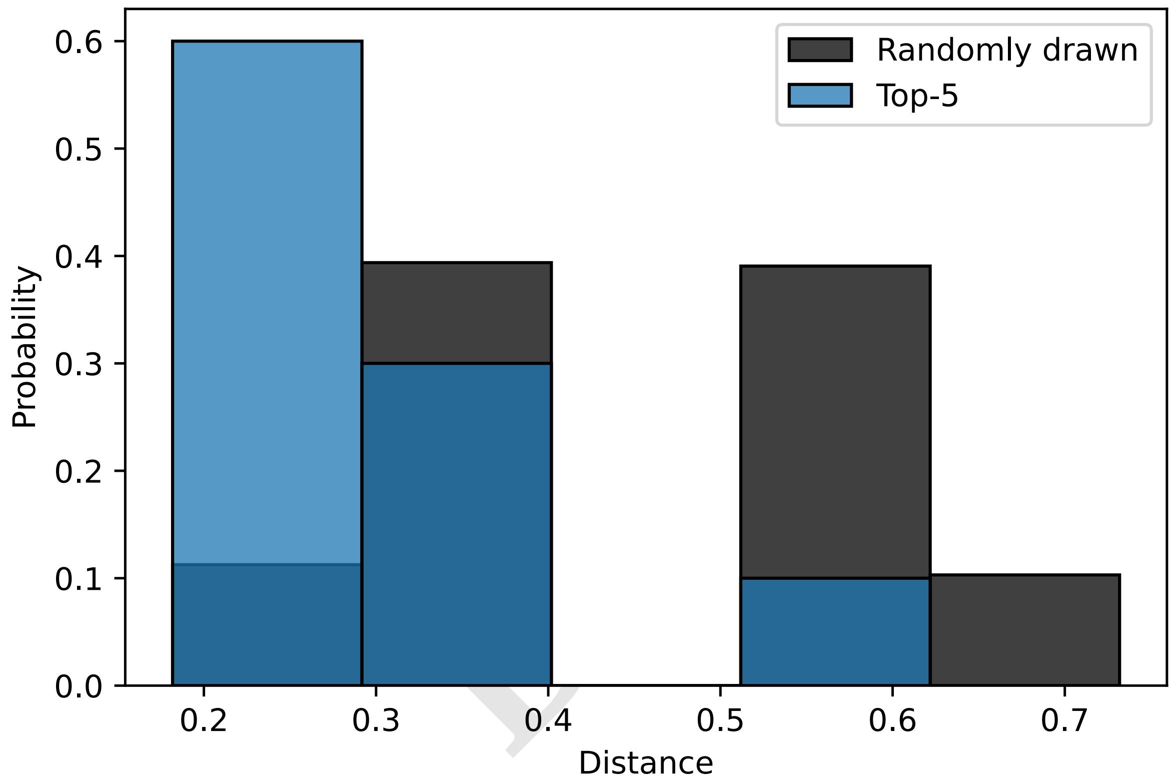


**Fig. S4.** Distribution of Hamming distance between all pairs of communities in our experiment (black) and between the pairs of the 5 communities with the highest average degradation scores (blue). The distributions are transparent and put on top of each other.

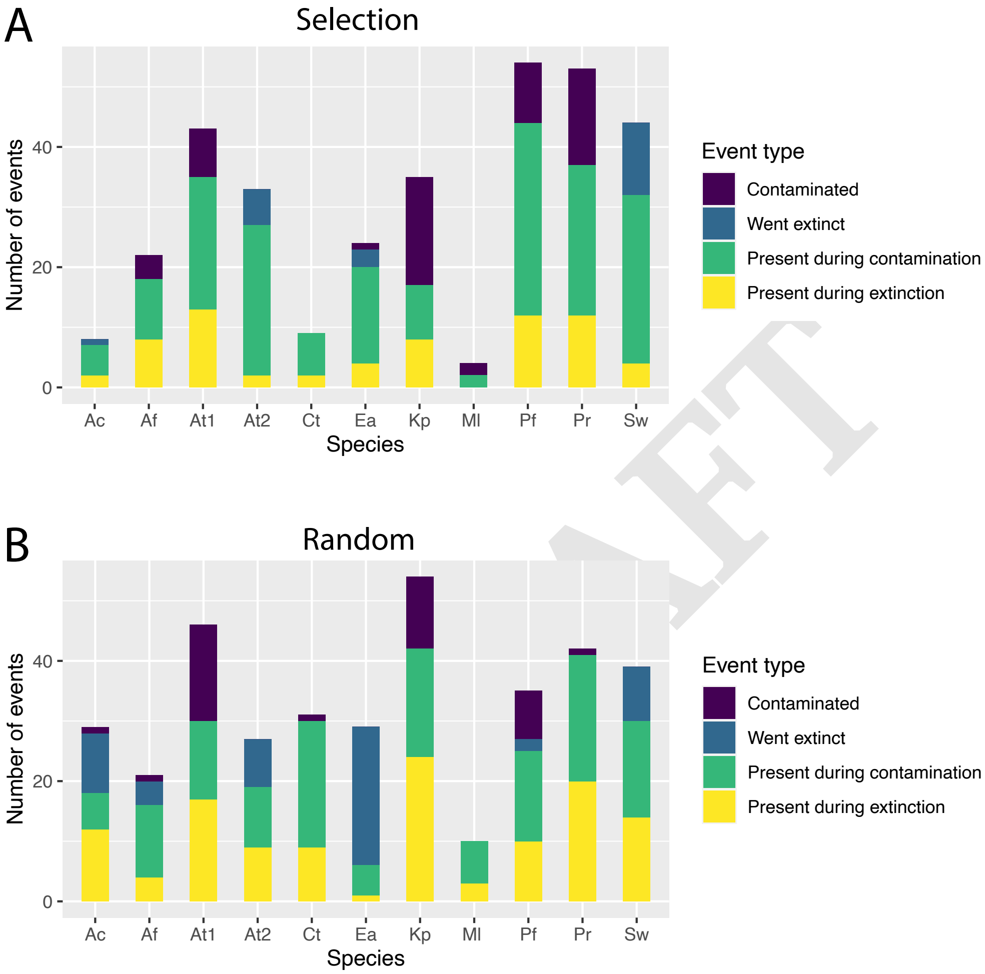


**Fig. S5.** Species' involvement in extinction and contamination events during the artificial selection experiment in A) the selection or B) the random treatment. We count the number of times a given species went extinct, contaminated a community, was present when an extinction occurred, or was present when a contamination occurred. Data over rounds shown in Tables S1.

DRAFT

| Round | Community composition | Extinct species |
|---|---|---|
| 0 | Sw_Kp_Pf_Pr | Sw |
| 1 | At1_Pf_Pr_At2 | At2 |
| 1 | At1_Pf_Pr_At2 | At2 |
| 1 | Sw_At1_Pf_At2 | At2 |
| 2 | At1_Pf_Pr_At2 | At2 |
| 2 | Sw_At1_Kp_Pf | Sw |
| 2 | Sw_At1_Ea_Pr | Ea |
| 3 | Sw_Kp_Pf_At2 | Sw |
| 3 | Sw_Af_Kp_Pf | Sw |
| 3 | Sw_At1_Pr_At2 | At2 |
| 4 | Sw_At1_Pr_At2 | Sw_At2 |
| 7 | Sw_At1_Ea_Pf | Sw |
| 8 | Sw_At1_Af_Ea | Sw |
| 9 | Sw_At1_Ea_Pr | Sw_Ea |
| 9 | Sw_At1_Af_Ea | Sw |
| 13 | Ea_Kp_Pf_At2 | Ea |
| 15 | Sw_Af_Ac_Kp* | Ac |
| 16 | Sw_Ct_Af_Ac | Sw |
| 16 | Sw_Ct_Af_Ac | Sw |
| 18 | Sw_At1_Kp_Pf | Sw |

| Round | Community composition | Extinct species |
|---|---|---|
| 0 | Sw_At1_Ea_Pr | Sw |
| 0 | At1_Ml_Pr_At2 | At2 |
| 1 | Sw_Ac_Ea_Kp* | Ea |
| 1 | At1_Ml_Pr_At2 | At2 |
| 1 | Sw_At1_Ac_Ea | Sw_Ea |
| 1 | Sw_Ac_Ea_At2* | Ea |
| 1 | At1_Ac_Ea_At2 | Ea |
| 2 | Sw_Ac_Kp_At2* | Ac |
| 2 | At1_Ac_Ea_At2 | Ea |
| 4 | Ct_Ea_Kp_Pf | Ea |
| 4 | Sw_Ac_Pr_At2* | At2 |
| 5 | Sw_Ac_Pr_At2* | Ac_At2 |
| 5 | Sw_Ac_Kp_Pr* | Ac |
| 5 | Ac_Ea_Kp_At2* | Ac_Ea |
| 6 | Sw_Ct_Kp_Pf | Pf |
| 6 | Sw_At1_Ea_Kp | Sw_Ea |
| 6 | At1_Ct_Kp_Pf | Pf |
| 6 | Sw_Af_Ac_Kp* | Ac |
| 7 | Sw_Ea_Pr_At2* | Ea_At2 |
| 8 | Sw_Ct_Ac_Kp | Sw |
| 8 | Sw_Ct_Af_Kp | Sw |
| 8 | Sw_Ea_Pr_At2* | Ea_At2 |
| 9 | Sw_At1_Ac_Kp | Sw |
| 9 | Ac_Ea_Pr_At2* | Ea_At2 |
| 10 | Ac_Ea_Pr_At2* | Ac_Ea |
| 10 | Ea_Pf_Pr_At2 | Ea |
| 10 | Af_Kp_Pf_Pr | Af |
| 11 | Ml_Ac_Kp_Pr* | Ac |
| 11 | Ea_Kp_Pf_Pr | Ea |
| 11 | Af_Ea_Kp_Pf | Ea |
| 11 | Ac_Ea_Pr_At2* | Ac_Ea_At2 |
| 13 | At1_Ac_Ea_Pr | Ea |
| 13 | Sw_Ct_Ac_Kp | Sw |
| 14 | Sw_Ct_Ea_Kp | Ea |
| 14 | Af_Ea_Kp_Pf | Ea |
| 15 | At1_Ct_Ea_Kp | Ea |
| 15 | Ea_Kp_Pf_Pr | Ea |
| 15 | Sw_Af_Kp_Pr* | Sw_Af |
| 16 | Sw_Af_Ac_Pr* | Af_Ac |
| 16 | Sw_At1_Pf_At2 | Sw |
| 16 | Ct_Ea_Kp_Pf | Ea |
| 17 | Sw_Af_Ac_Pr* | Af_Ac |
| 18 | Sw_Ac_Ea_At2* | Ea |

DRAFT

**Table S1.** Extinctions occurring in the selection (left) and random treatment (right). Asterisks mark communities that did not have any strong-growing species (At1, Ct, Pf), which tend to positively affect weaker growers (Fig. 4C) These communities may experience species extinction because they are missing these species, rather than through competition.

| Treatment | Type of interaction change | number |
|---|---|---|
| Random | Not significant in ancestor, now negative | 3 |
| Random | Negative in ancestor, now not significant | 2 |
| Random | Positive in ancestor, now not significant | 2 |
| Selected | Not significant in ancestor, now negative | 1 |
| Selected | Negative in ancestor, now not significant | 3 |
| Selected | Positive in ancestor, now not significant | 2 |

**Table S2.** Counts of interaction changes from Fig. 4 where we compare the ancestral to the evolved interactions. “Now” refers to the interactions between the isolates evolved under the respective treatment for 18 rounds. A total of 46 interactions were measured and could have changed in each treatment. Instead, we observe only 7 changes in the random treatment and 6 changes in the selection treatment.

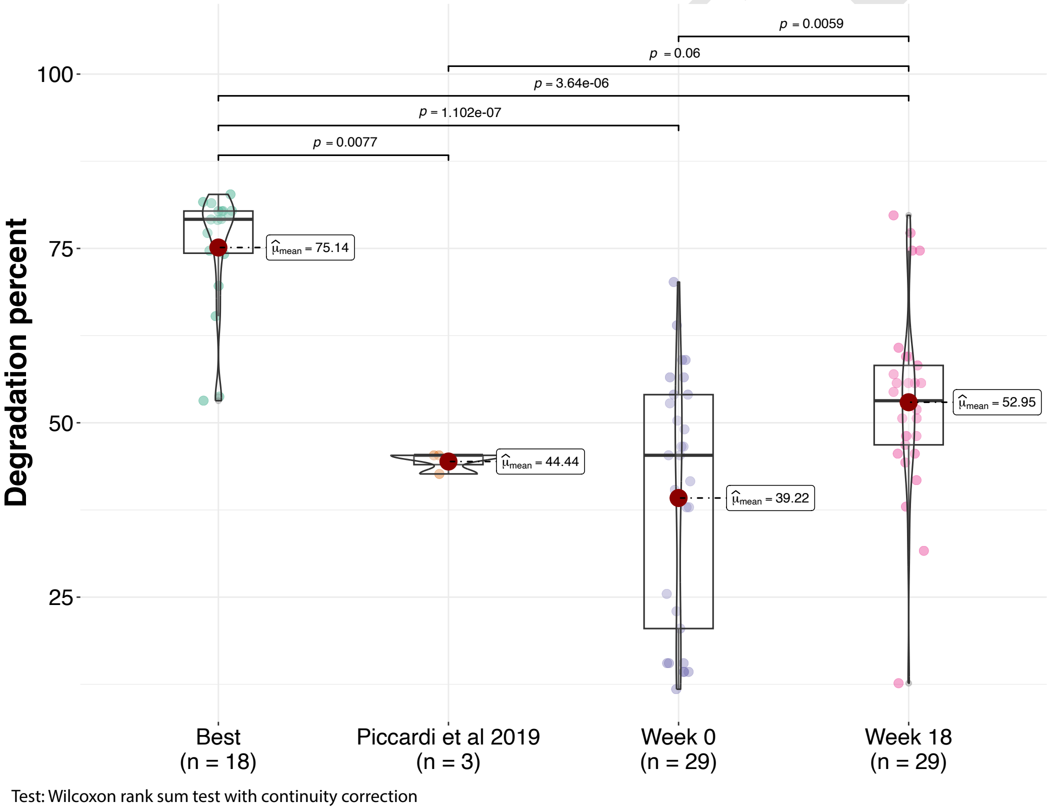


**Fig. S6.** Comparing the performance of the best community we found (“Best”) to the community studied in our previous work (“Piccardi et al 2019”). We also include the 29 communities in round 0 of the selection treatment and round 18. Statistical comparison used a Wilcoxon rank sum test.

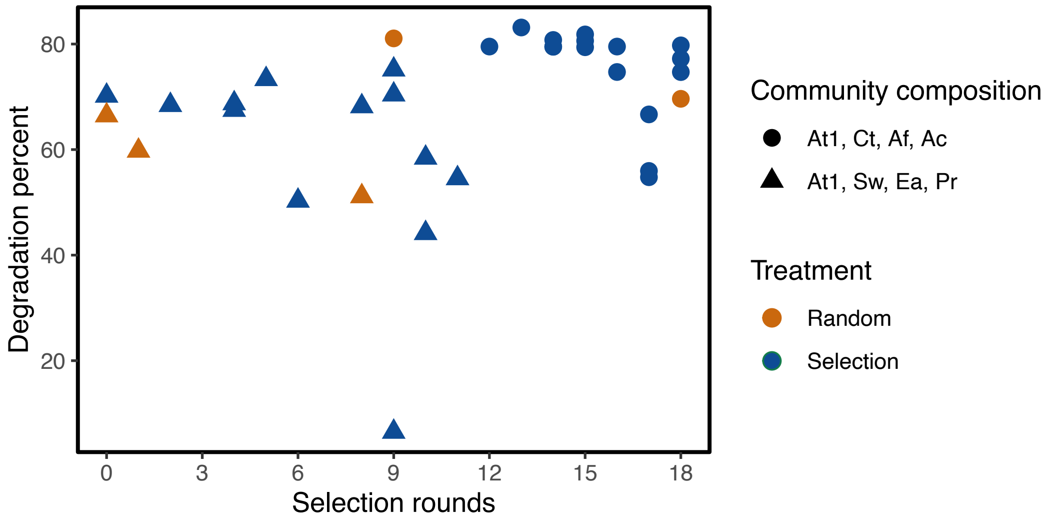


**Fig. S7.** Simulating how propagule selection would have performed: we plotted the degradation percent of the best-performing community at round 0 over all the rounds in which it appeared in both the selection and random treatments. We also plot all instances of the winning community with our approach. Neither of these communities improve in degradation score over time, but our winning community performs significantly better than the best community from round 0 (59.57 $\pm$ 16.83% versus 75.45 $\pm$ 8.43%, Wilcoxon p=0.0002).

| Species | Media name | Base medium | Base [g] for final vol. 300 ml | Antibiotic | Antibiotic conc. [µg/ml] |
|---|---|---|---|---|---|
| At1 | LB+KAN100 +SUL14.25+TRI0.75 | LB | 12 | KAN | 100 |
| | | | | SUL | 14.25 |
| | | | | TRI | 0.75 |
| Sw | MSA+NA15 | MSA | 33.3 | NA | 15 |
| Ct, At2 | Bru+SS2000+RIF50 | Brucella | 12.3 | SS | 2000 |
| | | | | RIF | 50 |
| Ml | NutrAgar+FO75 +AmB5+LEX8 +CST5+NA30 | Nutrient Agar | 8.4 | FO | 75 |
| | | | | AmB | 5 |
| | | | | LEX | 8 |
| | | | | CST | 5 |
| | | | | NA | 30 |
| Af | LB+STR200 +THY100 | LB | 12 | STR | 200 |
| | | | | THY | 100 |
| Ac | MMChromo+NA60 | MMChromo | 14.74 | NA | 60 |
| Ea | MRS+KAN25 | MRS | 18.6 | KAN | 25 |
| Kp | KIA+CAR100 | KIA | 12.24 | CAR | 100 |
| Pf | Cet+CAR50-Gly | Cetrimide | 14.01 | CAR | 50 |
| | | | no glycerol | | |
| Pr | PIA+TET15 | PIA | 13.5 | TET | 15 |
| | | | 6 ml glycerol | | |

**Table S3.** Recipes for selective media. All media were prepared to a total volume of 300ml. Abbreviations are as in Table S4.

DRAFT

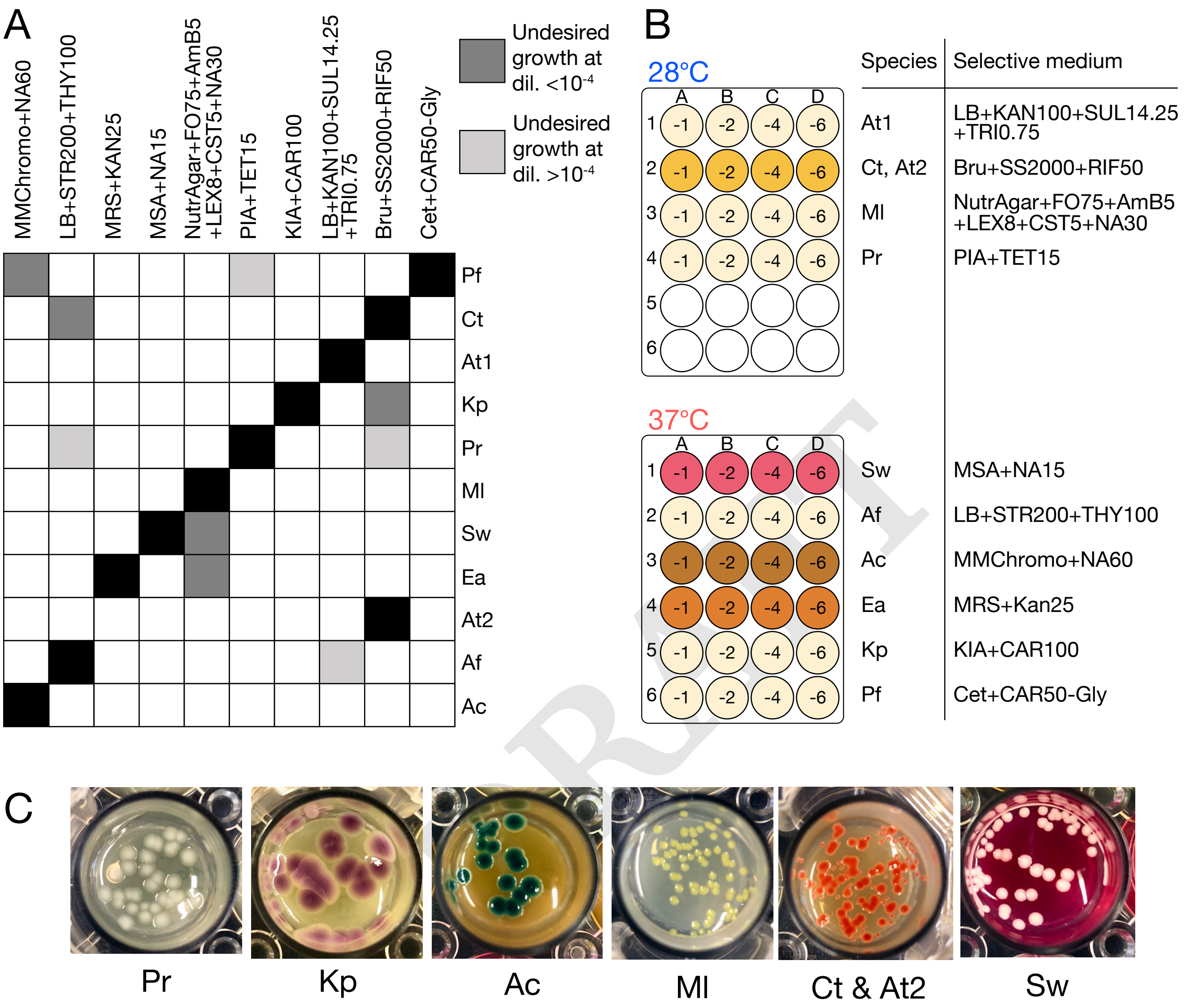


**Fig. S8.** Selective media for 11 species. (A) Black squares indicate no inhibition of a given species on the corresponding medium, dark grey squares indicate substantial growth (visible at dilution $10^{-4}$ or at higher dilutions), light grey indicates minor growth (visible at dilution $10^{-4}$ or at lower dilutions). We avoid combining species with dark grey squares but ignore the light grey ones and just avoid picking colonies from a low dilution. Nevertheless, this can explain some contamination patterns (Fig. S5). Abbreviations as in Table S4. Recipes for selective media are shown in Table S3. (B) 2×20 24-well plates were prepared in advance to contain 1.5ml per well of each selective medium in quadruplicate. We selected 20 communities to plate (10 from selection and 10 from random treatment), diluted each in 96-well plates containing PBS and inoculated 25 $\mu$l droplets of the corresponding dilutions into the wells as indicated above (dilutions -1, -2, -4 and -6). This gave us an approximate idea of the population sizes of each species in each of the 20 plated communities. (C) Six selective media with the seven species that could easily be distinguished on them growing in 24-well plates. As Ct and At2 could not be distinguished, we never combined them in the same community.

| Abbreviation | Full name |
|---|---|
| KIA | Klebsiella Isolation Agar (Sigma 90925) |
| MMChromo | Miller and Mallinson ChromoSelect Agar (Sigma 00563) |
| PIA | Pseudomonas Isolation Agar (BD 292710) |
| LB | Luria Bertani Agar (BD 244520) |
| Bru | Brucella Agar (Sigma 18795) |
| Cet | Cetrimide Agar (Sigma 22470) |
| NutrAgar | Nutrient Agar (Oxoid CM0003) |
| MRS | De Man, Rogosa, Sharpe Agar (Oxoid CM0361) |
| MSA | Mannitol Salt Agar (Oxoid CM0085) |
| RIF | Rifampicin |
| KAN | Kanamycin |
| SUL | Sulfamethoxazole |
| TRI | Trimethoprim |
| AmB | Amphotericin B |
| LEX | Cephalexin |
| CST | Colistin |
| NA | Nalidixic Acid |
| TET | Tetracyclin |
| FO | Fosfomycin |
| STR | Streptomycin |
| THY | Thymidine |
| CAR | Carbenicillin |
| SS | Sodium Selenite |

DRAFT

**Table S4.** Abbreviations used in medium ingredients and chemical sources.
